# Robust inference of historical human generation times

**DOI:** 10.1101/2023.07.20.549788

**Authors:** Richard J. Wang, Jeffrey Rogers, Matthew W. Hahn

## Abstract

Ragsdale and Thornton (2023) raise concerns about our recent estimates of historical human generation times, concluding that our results were “predominantly driven by nonbiological artifacts.” While we believe these authors have pointed out several important sources of uncertainty, we show here that their main concerns are either not relevant to our study or support our conclusions as much as they cast doubt on them. In particular, the demographic simulations carried out by Ragsdale and Thornton assume all individuals with recent African ancestry are from West Africa, which is not appropriate for our sample. In contrast to the lack of visual concordance between predictions and data cited by these authors as evidence for a lack of fit, we demonstrate that our model provides a good statistical fit to data on the overall historical mutation spectrum, though one particular mutation type is an outlier. Furthermore, we show that the historical generation times inferred when using alternative methods for estimating the ages of individual alleles are largely in agreement with our results, particularly so when using results from Relate. Importantly, these analyses, as well as recent work from an independent group, confirm the idea that a model built on *de novo* mutations and applied to polymorphism data provides useful and reliable estimates of generation times in widely distant mammals.

---

We thank Ragsdale and Thornton (2023) for their careful consideration of our recent study (Wang et al. 2023). These authors raise legitimate concerns about the uncertainty underlying our estimates of the human generation time, and present new data and analyses to consider. For example, they infer historical mutation spectra from two additional genealogical reconstruction methods, arguing that the resulting estimates of generation times (also called “generation intervals”) are not consistent with the ones we reported. Below, we address the issues raised in their paper, especially noting where we agree with them about the difficulties in estimating historical generation times. While these sources of uncertainty should certainly be considered, we also show that a statistical analysis of their new results provides further support for the robustness of our original conclusions.

## Ancestral population structure

Ragsdale and Thornton (2023) argue that our analyses require “long-lasting isolation among ancestral populations,” with population structure in humans stretching back 1-2 million years. This argument is based on the fact that our analyses show that the mutation spectrum differed in the ancestors of different human groups 10,000 generations ago and beyond. These historical mutation spectra rely on allele ages estimated by the program GEVA (Albers and McVean 2020). Our original paper noted the limited information on the mutation spectrum that could possibly be gleaned more than 10,000 generations into the past, which is why our analyses and discussion were limited to this interval. However, it may be our fault for including a figure-inset showing inferences beyond 10,000 generations—our intention was not to highlight these results, but instead to show that we were not hiding anything by using this cut-off. There are no error bars presented in this inset, so it is impossible to determine from it where the mutation spectra become indistinguishable among populations. We certainly do not make any inferences or claims about populations “many 10s of thousands of generations ago” in our paper.

Setting aside the issue of inferences beyond 10,000 generations ago (approximately 250,000 years ago), our results do clearly show differences in mutation spectra—and therefore generation times—in the ancestors of different human groups more recently than this point in time. In particular, our results imply that the ancestor of current samples with recent African ancestry (denoted AFR by the 1000 Genomes Project Consortium 2015) had a different mutation spectrum than the ancestor of samples with recent ancestry outside of Africa (denoted EAS, EUR, and SAS). As discussed in our paper, this result must reflect deep population structure within Africa, since all humans lived on this continent 250,000 years ago. Ragsdale and Thornton (2023) conclude that our inferences are incorrect, as even the deepest estimates from other studies put “the Eurasian-West African divergence at 100-150 ka [thousand years ago],” and most estimates put it closer to 75 ka. They carry out simulations to show how unreasonable it would be to have a signal of population structure between Europe and West Africa 250 ka, given a divergence time of 75 ka.

While we appreciate the detail of their simulation, it does not seem relevant to our results because it does not match our data. Although Ragsdale and Thornton (2023) continually refer to our sample as “West African,” it is not—the constituent sub-populations come from all over Africa and the African diaspora. In particular, the AFR continental sample we use includes the following population groups: Yoruba in Ibadan, Nigeria (YRI), Mende in Sierra Leone (MSL), Luhya in Webuye, Kenya (LWK), Gambian in Western Divisions in the Gambia (GWD), Esan in Nigeria (ESN), Americans of African Ancestry in South West USA (ASW), and African Caribbeans in Barbados (ACB). While several of these groups do currently live in West Africa, these samples reflect much more of the diversity of Africa, a continent on which recent work has inferred the existence of deep population structure more than 250,000 year ago (Fan et al. 2023; Pfennig et al. 2023; Ragsdale et al. 2023). Given this structure within Africa, and the fact that our sample is not exclusively West African, we do not think these results are inconsistent with the current understanding of human history. Finally, although it was not mentioned in Ragsdale and Thornton (2023), we note that the analyses they introduce using allele age estimates from the program tsdate also find this same difference in mutation spectra in Africa 250,000 years ago (their Figure S19; see next section for more detail on these results).

## Estimates of allele ages

As discussed above, Ragsdale and Thornton (2023) concluded that differences in mutation spectra among populations in Africa 250,000 years ago were incompatible with human history. To explain these results, they proposed that the allele ages inferred by GEVA are noisy and biased. If the ages of individual alleles provided by this method were faulty, then the resulting mutation spectra in each time period would be faulty, as would the generation times inferred from these spectra. As a first test of this idea, Ragsdale and Thornton (2023) compared the GEVA-inferred mutation spectra over time to the one predicted by our model. The inspiration behind this comparison is that the generation times predicted by our model themselves imply a mutation spectrum, and these predicted spectra can be compared to the spectra directly inferred from data as a test of model fit. Although we had previously presented an overall goodness-of-fit of our model (Figure S6 and S7 in Wang et al. 2023), the approach proposed by Ragsdale and Thornton has the advantage of examining the fit of each of the six mutation types on its own.

By comparing the GEVA-inferred mutation spectra over time (Figure 2A in Ragsdale and Thornton 2023) to the spectra predicted by our model (Figure 2D in Ragsdale and Thornton 2023), the authors concluded that “the inferred generation times provide a poor fit to the data.” We agree that the visual match between these two plots seems poor. However, no further investigation of the data is presented by Ragsdale and Thornton beyond the seeming lack of visual concordance. We wondered whether a statistical analysis—or a different graphical representation—might reveal something further about the fit of our model to the data.

Figure 1 shows the individual predictions and data for the six different mutation types. This view of the results makes it much easier to appreciate where our model fits the data well and where it does not. As can be seen, for three of the six mutation types the statistical fit is very good (A→ T, C→A, and C→G all have *R*^2^>0.7; Figure 1), for two mutation types it is fairly good (A→C and C→T have *R*^2^>0.3), and for one it is poor (A→G has *R*^2^≈0). While there is clearly substantial variance among mutation types in how well our model fits, we think the overall fit is quite impressive, especially for a model that predicts the mutation spectrum based solely on changes in the generation time. Interestingly, a recent paper (Beichman et al. 2023) examining the mutation spectrum across multiple mammals also found that A→G mutations were not well-predicted by the same *de novo* mutation data we used to parameterize our model. Further work is clearly needed to understand why this mutation type behaves in this manner.

**Figure 1.**
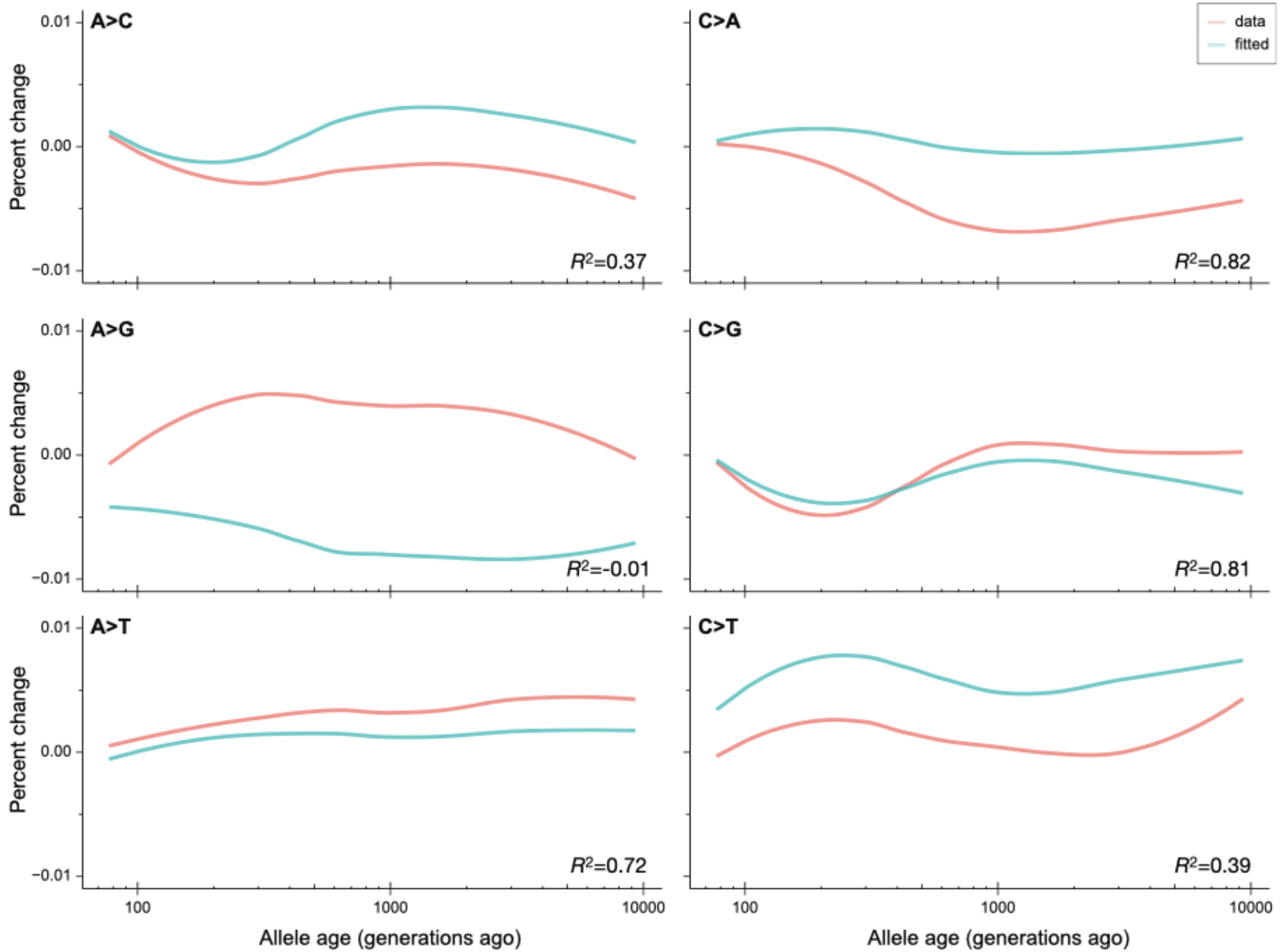
Observed (“data”) and estimated (“fitted”) percent change in the frequency of each mutation class through time, anchored to the most recent time window. The data here are generated by GEVA, with spectra estimated using the Dirichlet-multinomial model presented in Wang et al. (2023). The coefficient of determination (*R*^2^) between observed and estimated change in frequency through time is also shown for each mutation class.

As a second test of the idea that GEVA provided biased and noisy data, Ragsdale and Thornton (2023) use two alternative methods for dating allele ages: tsdate (Wohns et al. 2022) and Relate (Spiedel et al. 2019). Although we did not assume that the point estimates of allele ages from GEVA were necessarily correct—we also showed that we obtained very similar results when sampling ages from the posterior of GEVA estimates (Figure S14 in Wang et al. 2023)—it is always good to see how robust a result is to the choice of data and software. Both tsdate and Relate performed well in a recent evaluation (Brandt et al. 2022), though unfortunately GEVA was not included in that comparison.

After estimating historical mutation spectra from tsdate and Relate, Ragsdale and Thornton (2023) used our model to predict generation times with each dataset. They conclude that analyses using the data from these methods “provide qualitatively different inferred generation time histories.” There is no further comparison of the generation time histories inferred using tsdate and Relate (though they do compare the datasets themselves), and the histories themselves are only shown in the supplementary materials (Figures S18 and S19). We used these results in order to carry out a statistical analysis and to explore differences and similarities with our original predictions.

Here, we plot the predicted generation times for males and females using the original GEVA-based data (Figure 2A) beside those from tsdate (Figure 2B) and Relate (Figure 2C). There are clear differences between the three sets of predictions, but also striking similarities. For instance, all methods predict longer male generation times across the entire period, a higher variance in male generation times compared to female generation times, as well as a decrease in generation times from approximately 1,000 generations ago to 200 generations ago. Interestingly, there is a much higher correlation between our original predictions and those from Relate for males alone, females alone, as well as sex-averaged generation times (Table 1). Consistent with this observation, the average male and female generation times estimated using Relate (32.0 and 25.9 years, respectively) are within the standard errors of our original estimates using GEVA (30.7 ± 4.8 years for males and 23.2 ± 3.0 years for females). As can be seen in Figure 2, estimates from tsdate are almost always higher (37.1 and 26.5 years for males and females, respectively). While there are still important open questions as to which method provides the most accurate allele ages (if there is just one model appropriate for all mutation types), we think that the quantitatively and qualitatively similar results among methods speaks to the robustness of our original conclusions, rather than any problems unique to them.

**Figure 2.**
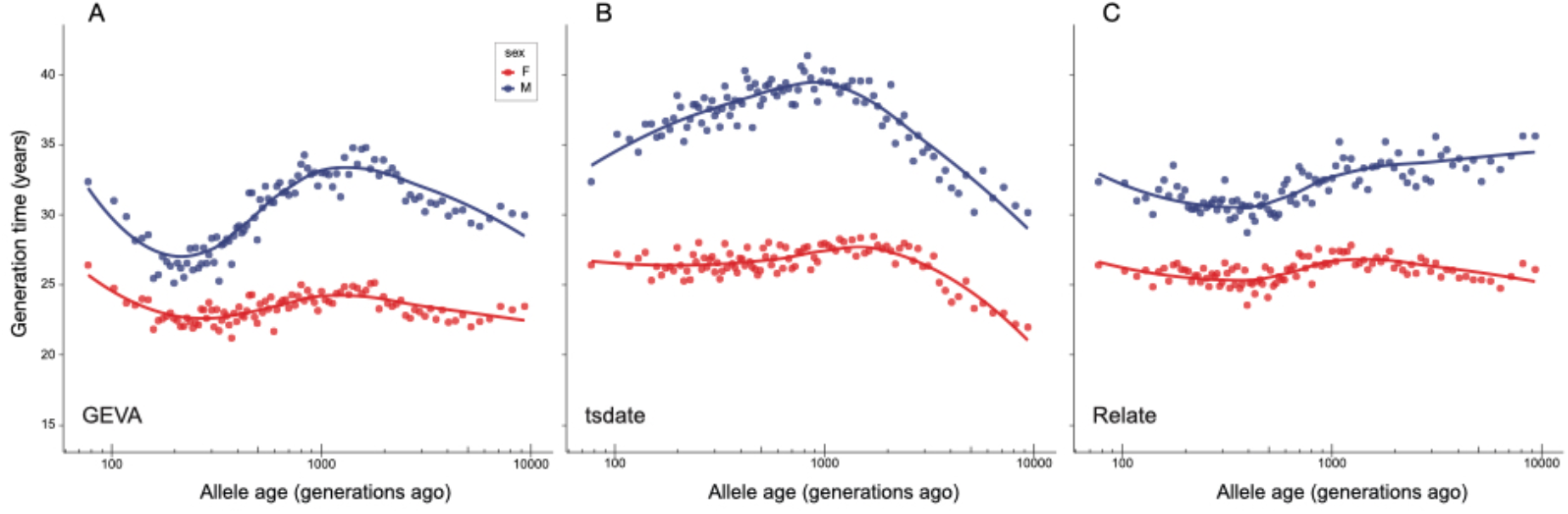
Generation times estimated for males and females across the past 10,000 generations. Estimated were generated from three different datasets: A) GEVA, B) tsdate, and C) Relate.

**Table 1.**
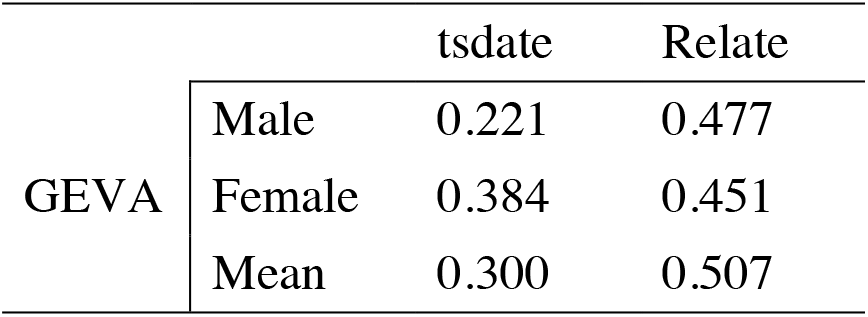
Pearson correlation (*r*) between GEVA-based estimates of generation times and those using tsdate and Relate. Estimates from the past 10,000 generations were used.

| GEVA |  | tsdate | Relate |
| --- | --- | --- | --- |
|  | Male | 0.221 | 0.477 |
|  | Female | 0.384 | 0.451 |
|  | Mean | 0.300 | 0.507 |

## Spectra of *de novo* mutations and polymorphisms

Ragsdale & Thornton (2023) note that in our original paper we reported a difference between the *de novo* mutation spectrum from Icelandic trios (Jónsson et al. 2017) and the spectrum from the youngest bin of polymorphisms (Table S1 in Wang et al. 2023). We discussed this difference in our paper—though we were unable to uncover its source—and proposed a statistical method for estimating generation times despite this discrepancy (and regardless of its cause). We tested some of the assumptions of this method and obtained similar results (see section S4.4 of Wang et al. 2023). Our paper acknowledges that we are not able to estimate reasonable generation times without this correction for the difference in spectra.

The assumptions of the correction are clearly stated in our paper, and Ragsdale and Thornton (2023) are of course not obliged to agree with them. Their paper explores some possible explanations for the discrepancy in mutation spectra, but we disagree that a full accounting for this difference is necessary to correct for it. We proposed a statistical correction using transparent and appropriate methods, and none of the results presented by these authors establish that this correction is invalid or incorrect. For instance, we do not think that the disagreement between one high-quality estimate and one low-quality estimate of the *de novo* mutation spectrum in humans is evidence that the data are of overall low quality or are affected by bioinformatic errors. We note that we also obtained highly similar results using the lower quality *de novo* dataset (see Figure S12A in Wang et al. 2023). Regardless, we agree that it will be informative going forward to understand the source of the discrepancy, especially as a difference between the spectrum of *de novo* mutations and polymorphisms may be common across species (e.g. Schrider et al. 2013; Zhu et al. 2014; Carlson et al. 2018; Wang et al. 2022; Beichman et al. 2023).

## Conclusions

It is difficult to estimate historical generation times from polymorphism data. One needs both a well-parameterized model of how the mutation spectrum changes with parental age and accurately dated ages of polymorphic alleles. There are many sources of uncertainty and error in both of these tasks, and it is understandable that Ragsdale and Thornton (2023) would want to take a closer look at how this was done in our study. While their paper raises important questions and contributes new analyses and datasets, the take-home message of the further analyses presented here is that our original results provided a good statistical fit to the data and were largely robust to the methods being used. We are also reassured by the fact that similar models using the *de novo* mutation spectrum in humans are good fits to data from diverse mammals (Wang et al. 2022; Beichman et al. 2023). Nevertheless, we look forward to more sophisticated models that build upon and improve these results, thereby providing even more accurate inferences of historical generation times.

